# ClearFinder: a Python GUI for annotating cells in cleared mouse brain

**DOI:** 10.1101/2024.06.21.599877

**Authors:** Stefan Pastore, Philipp Hillenbrand, Nils Molnar, Irina Kovlyagina, Monika Chanu Chongtham, Stanislav Sys, Beat Lutz, Margarita Tevosian, Susanne Gerber

**Affiliations:** Institute for Human Genetics, University Medical Center Johannes Gutenberg University, 55131 Mainz, Germany; Institute of Pharmaceutical and Biomedical Sciences, Johannes Gutenberg University, 55128 Mainz, Germany; Institute of Physiological Chemistry, University Medical Center Johannes Gutenberg University, 55128 Mainz, Germany; Leibniz Institute for Resilience Research; 55122 Mainz, Germany

**Keywords:** tissue clearing, atlas alignment, cell count, 3D volumetric imaging

## Abstract

**Background:** Tissue clearing combined with light-sheet microscopy is gaining popularity among neuroscientists interested in unbiased assessment of their samples in 3D volume. However, the analysis of such data remains a challenge. ClearMap and CellFinder are tools for analyzing neuronal activity maps in an intact volume of cleared mouse brains. However, these tools lack a user interface, restricting accessibility primarily to scientists proficient in advanced Python programming. The application presented here aims to bridge this gap and make data analysis accessible to a wider scientific community.

**Results:** We developed an easy-to-adopt graphical user interface for cell quantification and group analysis of whole-cleared adult mouse brains. Fundamental statistical analysis, such as PCA and box plots, and additional visualization features allow for quick data evaluation and quality checks. Furthermore, we report significant differences in total cell counts between CellFinder and ClearMap when cross-analyzing the same samples, underscoring the need for optimizing reproducibility within the field.

**Conclusions:** Our easily accessible tool allows more researchers to implement the methodology, troubleshoot arising issues, and develop quality checks, benchmarking, and standardized analysis pipelines for cell detection and region annotation in whole volumes of cleared brains.

## Background

Immediate-early gene (IEG) expression has been used for decades to visualize populations of activated neurons. Nowadays, activity-dependent promoters and transgenic reporter mouse lines, allowing for permanent labelling of neurons at selected time points, are widely implemented by neuroscientists [1]. However, to detect fluorescently tagged ensembles of neurons distributed throughout the brain, the tissue has to be sectioned, resulting in bias and artifacts. A new approach to overcome these limitations was recently developed: tissue clearing combined with light-sheet fluorescence microscopy. This concept gained popularity among neuroscientists studying the complexity of the nervous system [2].

One of the major applications of this methodology is to identify the brain regions where neuronal activity is down- or upregulated. This is achieved by first detecting individual cells in a 3D volume and then performing a non-linear transformation of the images to the reference atlas. However, the acquisition of the complete volume of a cleared adult mouse brain can result in several terabytes of data, necessitating a sophisticated infrastructure with high data storage capacity and powerful computational resources for processing.

Currently, ClearMap [3,4] and CellFinder [5] are the only open-source tools for cell detection and annotation in whole/intact brain volume. However, both software solutions - albeit very useful - are command-line based and require third-party packages that are frequently not maintained. In addition, neither tool offers statistical analysis and data visualization functionality. Despite some attempts at making these pipelines more accessible to biologists and microscopists [6], the installation and usage still requires intensive support from skilled bioinformaticians or computer scientists.

Here, we developed ClearFinder, a Graphical User Interface (GUI) unifying the functionality of both ClearMap and CellFinder, while at the same time simplifying the setup process and offering fundamental statistical analysis of results. The user is guided through streamlined data selection and preparation steps, including uniform file renaming, adjustment of cell detection parameters and image pre-processing, alignment to a reference atlas, and generation of an output file containing cell counts per brain region.

## Materials and Methods

### Sample preparation

All experiments were performed according to the European Community’s Council Directive of 22 September 2010 (2010/63EU) and approved by the respective agency of the State Rhineland-Palatinate (Landesuntersuchungsamt, permit number G-17-1-021). Male mice were group-housed in temperature- and humidity-controlled rooms with a 12-hour light-dark cycle with water and food provided ad libitum. Seven days prior to behavioral experiments, animals were single-housed. Mice used in this study were eight weeks old by the start of the experiments. B6.Cg-Tg(Arc-cre/ERT2)MRhn/CdnyJ mice (JAX Nr. 022357 [7]) were crossed with B6.129-Gt(ROSA)26Sortm5(CAG-Sun1/sfGFP)Nat/J mice [8], and generated double transgenic heterozygous mice are referred to as Arc-nuclGFP for simplicity. Upon neuronal activity, in the presence of tamoxifen (Merck, Germany), neurons in these mice can express a nuclear membrane variant of green fluorescent protein (Sun1/sfGFP), thereby obtaining permanent labelling of these cells. Mice received an intraperitoneal (i.p.) injection of tamoxifen (150 mg/kg) during a stressful behavior paradigm [9]. Afterwards, mice were deeply anesthetized by i.p. injection of 100 mg/kg pentobarbital (Merck, Germany) and 0.1 mg/kg buprenorphine (TEMGESIC, Indivior Europe Ltd, Ireland) in water. After checking for the absence of the interdigital reflex, mice were perfused transcardially with 100 ml ice-cold phosphate-buffered saline (PBS), pH 7.8, and 50 ml 4% freshly prepared paraformaldehyde (PFA) solution. The brain was dissected and post-fixed in 4% PFA at 4°C for 8 hours, then washed in PBS 3 times for 30 minutes each.

iDISCO+ tissue clearing was performed according to Jin et al. [10] with modifications. Briefly, dissected tissue was washed in PBS and dehydrated at room temperature (RT) in MeOH (Merck, Germany) solutions in water: 20, 40, 60, 80, 100% MeOH for 1 hour each, followed by incubating in 100% MeOH overnight at RT. Afterwards, tissue delipidation was performed by incubating the samples in dichloromethane (DCM; Merck, Germany) and MeOH solution (1:3) twice for 2 hours at RT. After washing the samples for 1 hour in 100% MeOH, they were transferred to 5% H_2_O_2_ (Merck, Germany) in 100% MeOH solution overnight at 4°C. After rehydrating the samples in decreasing concentrations of MeOH solution in water at RT (80, 60, 40, 20% MeOH for 1 hour each), they were transferred to permeabilization solution (78.6% PTx0.5 (PBS with 0.5% Tween 20 and 10 mg/ml heparin); 1.4% glycine; 20% dimethyl sulfoxide, DMSO) for 2 days at 37°C, followed by blocking solution (84% PTx0.5 (PBS with 0.5% TritonX-100); 6% normal donkey serum, NDS; 10% DMSO) for 2 days at 37°C. Afterwards, the samples were incubated at 37°C in 1:1000 diluted primary antibody (chicken anti-GFP, Aves Labs, Davies, CA, USA; RRID:AB_10000240) solution (92% PTwH0.5; 3% NDS; 5% DMSO) for 7 days, with sequential additions of the primary antibody on days 2, 3, 4 and 5, thereby reaching a final dilution of 1:200. The samples were washed in PTwH0.5 over 4 days at 37°C and then incubated with the 1:500 diluted secondary antibody (Alexa Fluor® 647 AffiniPure F(ab’)_2_ Fragment Donkey Anti-Chicken IgG (H+L), Jackson ImmunoResearch, West Grove, PA, USA) solution (92% PTwH0.5; 3% NDS; 5% DMSO) for 7 days at 37°C with sequential additions of the antibody on days 2, 3, 4 and 5, thereby reaching a final dilution of 1:100. The samples were washed in PTwH0.5 over 4 days at 37°C. Then, the samples were dehydrated and delipidated again, as described above. Afterwards, they were incubated in dibenzyl ether (DBE; Sigma-Aldrich, Germany) overnight at RT for refractive index (RI) matching until fully transparent. Samples were stored and imaged in DBE at RT. All incubation procedures were performed under constant gentle mixing on a nutating mixer (ThermoFisher Scientific, USA)

Microscopy of cleared samples was performed in horizontal orientation on the light-sheet microscope UltraMicroscope II (LaVision Biotec) with a camera (Andor Neo sCMOS) and a 2x/0.5NA objective (MV PLAPO 2XC, Olympus) with corrected dipping cap attached. Exposure time was set to 150 ms, sheet width to 100%, zoom factor 1.0x, light-sheet merging algorithm – blend. Acquired images were processed using Vision4D (Carl Zeiss Microscopy Software Center Rostock GmbH, Germany).

### System requirements

Recommended PC requirements

OS: Linux (Debian Ubuntu)

CPU: Intel(R) Xeon(R) Silver 4210R CPU @ 2.40GHz

Processors: 20 (Threats)

RAM: 128GB

## Implementation

ClearFinder is written in Python using the QT framework. The integration of Napari [11], CellFinder, and ClearMap environments management, as well as job scheduling is orchestrated by Nextflow [12]. The workflow (Fig. 1) is directly applicable after the collection of TIFF files from stitched images from the light-sheet microscopy experiment. Detected cells of ClearMap are converted into XML format, enabling a CellFinder-like cell visualization in Napari. The ClearFinder GUI additionally allows the training of CellFinder’s neuronal network for cell classification. Training data can be obtained with a CellFinder plugin for Napari. Detected cells are embedded into a comparable data structure. The latter allows a summary of cell counts over different structural hierarchies based on the phylogeny of the Allen Brain Atlas (http://alleninstitute.org/) mouse ontology file. Cell counts of several samples can be integrated into a single data frame with an additional option to create a metadata frame, allocating the samples to respective experimental groups.

**Figure 1.**
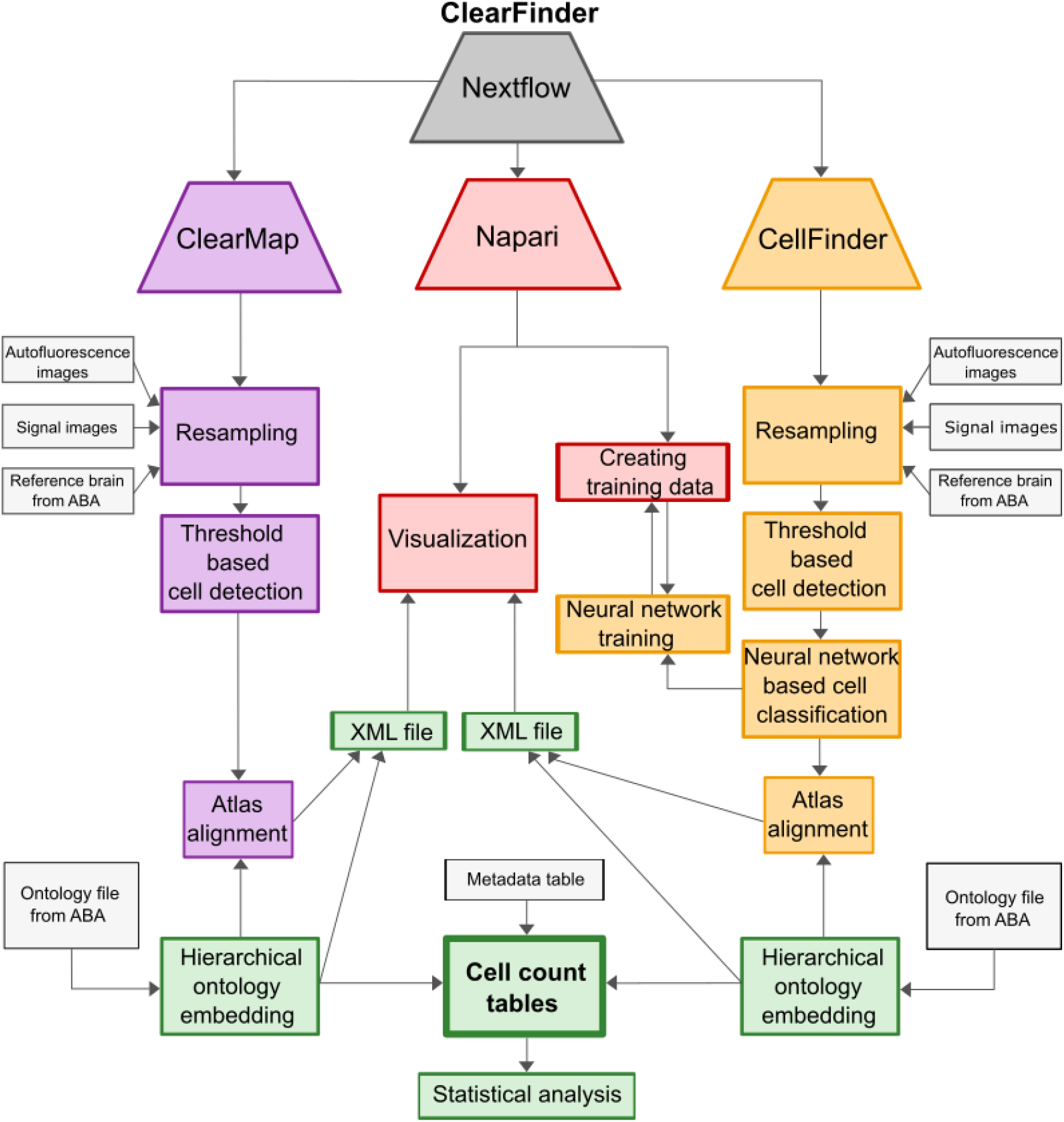
ClearFinder GUI architecture. Schematic representation of the workflow, unifying both analysis tools under one GUI. Trapezoids indicate individual software packages: violet – ClearMap, red – Napari, orange – CellFinder, grey – Nextflow. Separate analysis steps with each package are indicated in rectangular boxes of respective colors. Light-grey boxes indicate the input from the user. Green boxes indicate the output of ClearFinder GUI. ABA – Allen Brain Atlas.

Furthermore, two normalization options for the number of cells detected across different samples and experimental conditions are available: median of ratios [13] or cells per million (Fig. 2). A unified data output format allows a standardized statistical analysis between conditions across both tools. CellFinder, ClearMap, and Napari are embedded in independent Conda environments and are available on GitHub (https://github.com/stegiopast/ClearFinder). Anaconda environments are an essential feature of the overall software design due to their practical ability to manage Python environments while simultaneously allowing the installation of required non-Python dependencies, such as CUDA, which are difficult to manage manually. Furthermore, placing CellFinder, Napari, and ClearMap in different environments allows the integration of tools into one whole software suite while keeping dependencies. Overall, this strategy guarantees future stability.

**Figure 2.**
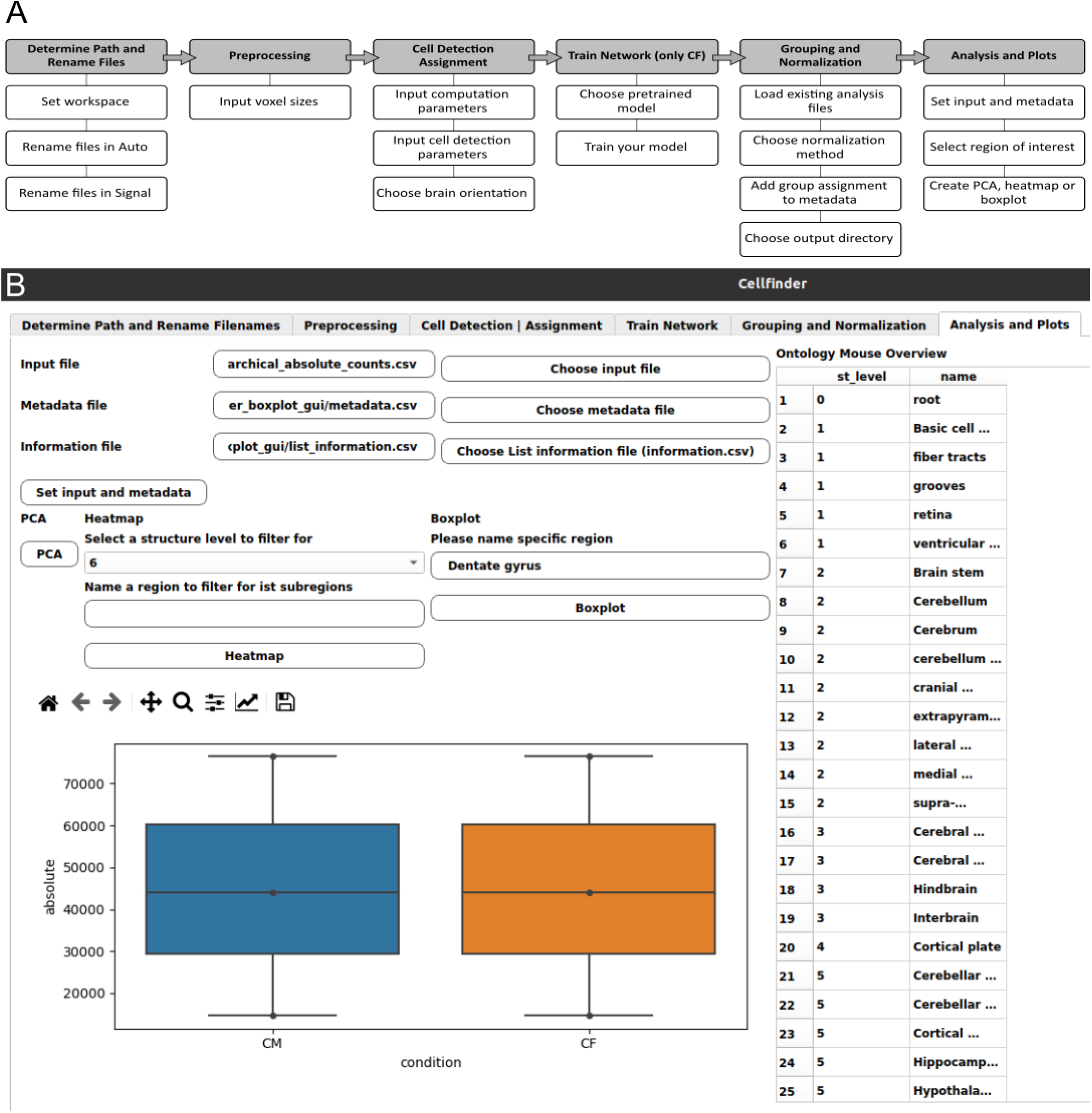
ClearFinder GUI interface. **A:** A wizard illustrating a step-by-step process of loading, processing and analysis of the data, orchestrated by the ClearFinder GUI. Top row represents the major processing steps, including the selection options in black boxes below. **B:** A screenshot of the GUI interface depicting the last step of the GUI wizard allowing basic analysis and visualization of obtained results.

## Results

### ClearFinder GUI for neuronal activity mapping

Although both ClearMap and CellFinder are potent tools for cell detection in cleared tissue [14–18], both programs are quite challenging in terms of installation, data handling, setting the parameters for cell detection, and image transformation. These steps require manual execution via the command line.

To address these limitations, we developed a unified, intuitive interface: ClearFinder (Fig. 1-2). The GUI offers several advantages, including streamlined setup for Ubuntu-based systems, renaming images and setting workspaces to process the gathered data. Subsequently, processing parameters for the resampling and cell detection steps can be adjusted easily (Fig. 2).

Moreover, ClearFinder integrates both ClearMap and CellFinder in a single toolbox by a Nextflow script workflow (Fig 1). As a result, users can initiate both tools with a single bash command. Following independent processing with ClearMap and CellFinder, the generated results can be easily visualized and compared in Napari, providing a cohesive data output for subsequent in-depth statistical analysis.

### ClearFinder reveals differences in total cell counts between ClearMap and CellFinder

To compare the performance of cell detection and alignment in CellFinder and ClearMap, we analyzed one dataset (n=3) with ClearFinder. To this end, Arc-nuclGFP [7,8] (see Materials and Methods) mice were injected with tamoxifen after a chronic stress exposure to induce permanent GFP labeling of neuronal nuclei throughout the brain. Later, the brain tissue was dissected, stained, cleared using iDISCO+ and imaged on a light-sheet microscope (Ultramicroscope II) in whole volume. After initial preprocessing (tile scan stitching using commercial software arivis Vision 4D), the data was analyzed using ClearFinder. We performed cell detection with three different size-thresholds (4, 5, and 6 μm in three axes), then the cells were allocated to the mouse Allen Brain Atlas [19]. The principal component analysis (PCA, Figure 3A) conducted on the total number of GFP-positive cells across all brain regions unveiled considerable disparities between the methods and thresholds. Notably, we detected high variability among samples subjected to cross-analysis using identical thresholds but different methods. In a 2-way ANOVA, total cell counts significantly differed between detection methods, but not thresholds (Figure 3B). Since the threshold value of 555 (5 μm in each axis) produced results with the lowest within-group variability for both methods, we selected it for further analysis.

**Figure 3.**
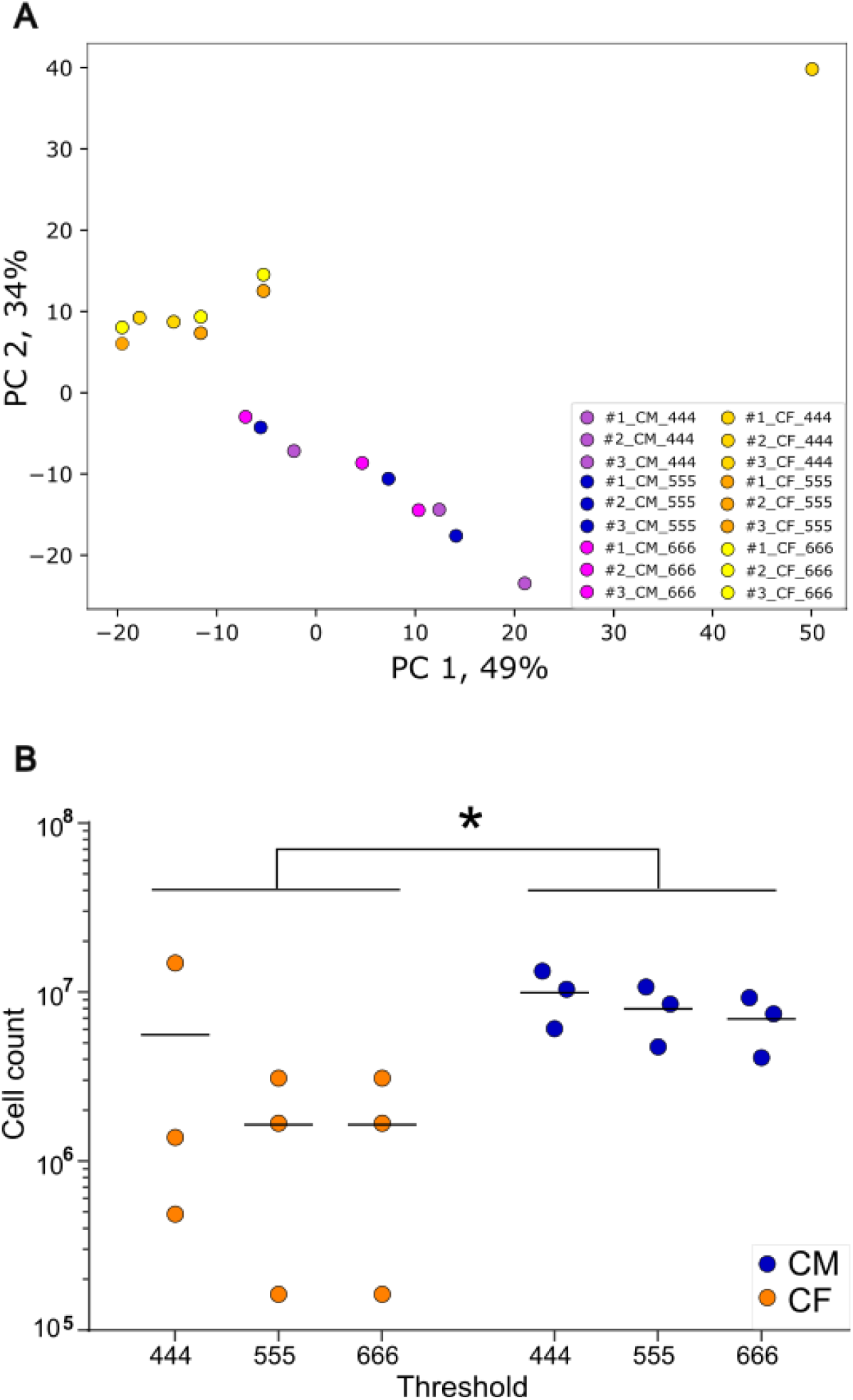
Differences in total cell counts between ClearMap and CellFinder. PCA plot of all cell counts in/across all brain regions. GFP-positive cells were detected in all regions of three sample using two methods and three detection thresholds. Yellow, orange and dark-orange dots indicate data obtained with CellFinder (CF), magenta, blue and violet dots indicate data processed with ClearMap (CM). CF dots were displaced by 2% vertically to avoid a complete overlap. PC – principal component. **B**, Total cell counts across all regions. Total counts of GFP-positive cells detected across all brain regions using two methods and three detection thresholds. The y axis represents the log-transformed cell counts. Significant differences between methods identified by 2-way ANOVA, pmethod = 0.0165, pthreshold = 0.3131, pmethod x threshold = 0.3131. * p < 0.05. CF – CellFinder, CM – ClearMap.

### ClearFinder reveals discrepancies in cell detection efficiency

The brain regions annotated in the Allen Brain Atlas are organized in 12 ontological levels from 0 being the root, containing all other brain regions, to 12, encapsuling the smallest subregions and nuclei. ClearFinder allows the sub-selection of ontological levels for easy analysis of a region of interest. We selected the hippocampal region (ontological level 6) for a side-by-side visual inspection of cell detection. The cells were detected using CellFinder (Figure 4 A, C) and ClearMap (Figure 4 B, D), and visualized in Napari. The comparison further highlighted noticeable differences in cell detection between the two tools. Noteworthy, the performance of CellFinder was not consistent. CellFinder detected fewer cells in sample #3, compared to ClearMap, while comparable number of cells were detected in sample #2. These prominent differences cannot be attributed to the additional machine learning-based cell/non-cell discrimination, since the number of non-cells was negligible in both analyzed samples (blue circles in Figure 4 A-D). Additionally, cell/non-cell discrimination functionality is absent in ClearMap. Further, we visualized cell counts across the subregions of the hippocampal area (Figure 4E), as well as a selection of cortical (Suppl. Fig. 1) and subcortical regions (Suppl. Fig. 2) as heatmaps. We observed consistently higher cell counts in all samples analyzed with ClearMap than with CellFinder.

**Figure 4.**
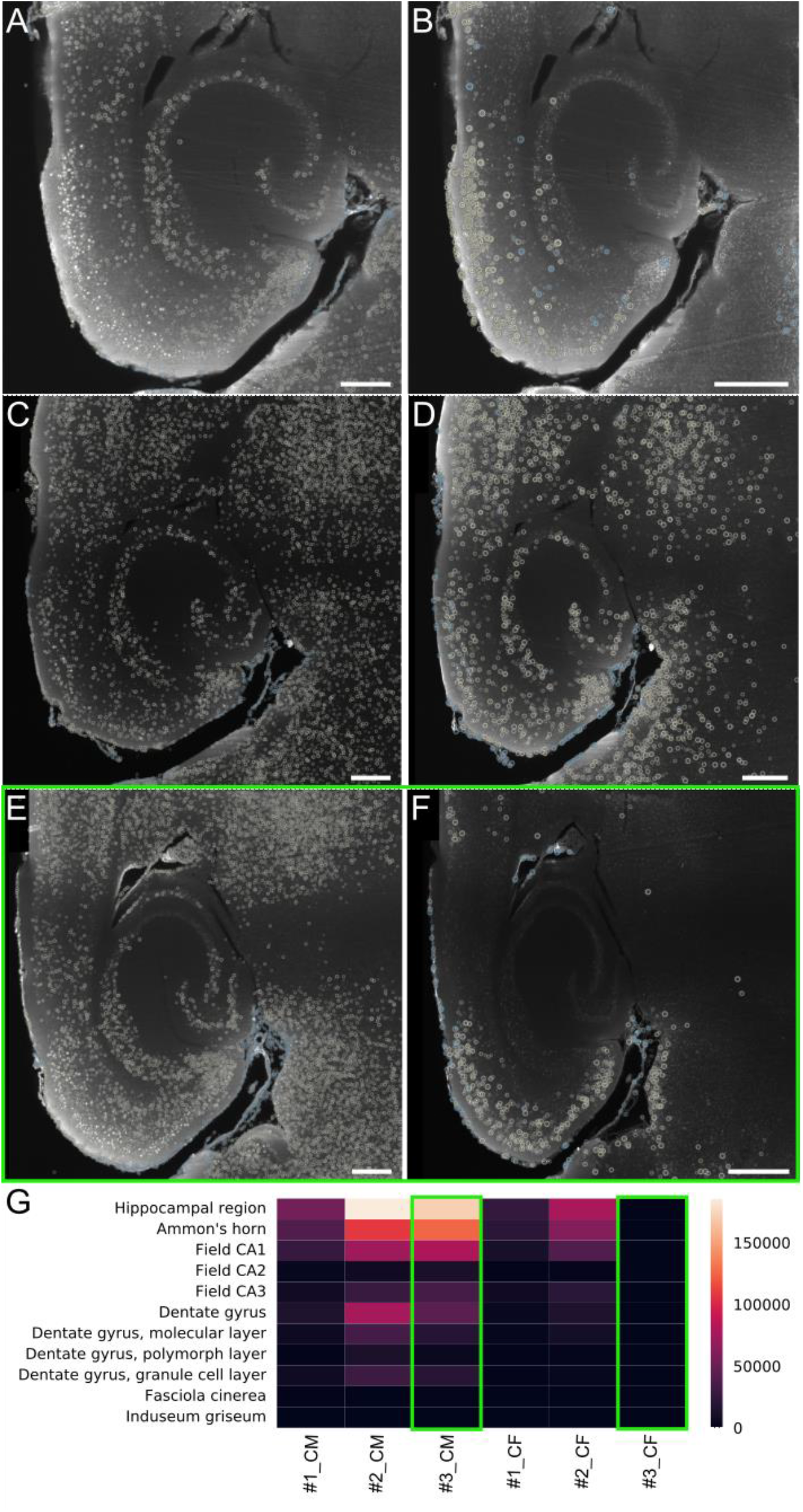
Cell detection efficiency. Sample #1 and Sample #2 analyzed with ClearMap (**A** and **C**) and CellFinder (**B** and **D**) using threshold 555 Sample #3 analyzed with ClearMap (**E**) and CellFinder (**F**) using threshold 555, marked by a green rectangle. Yellow circles indicate cell maxima: “cells”. Blue circles indicate maxima unassigned to a brain region: “non-cells”. The radius of the cells indicates the proximity to the cell core in the selected optical plane. Smaller circles indicate the cell maxima out of plane. Scale bar: 150 μm. **E**: Heatmap indicating the number of cells detected in subregions of the hippocampal area, as detected in each sample by ClearMap (CM) and CellFinder (CF) using the same threshold. Sample #3 marked by a green rectangle. Color-bar corresponds to cell count.

In summary, ClearFinder facilitates cell detection in cleared mouse brains with both ClearMap and CellFinder in a unified graphic-user-assisted pipeline. Integrated features also enable visualization and statistical analysis of results. Side-by-side comparison of cell counts obtained with both ClearMap and CellFinder also allows users to focus on samples with consistent yield irrespective of the detection strategy. This validation process circumvents biases due to data processing and thereby enhances the robustness and biological validity of the findings.

### Documentation

An extensive documentation and the code are available on the project’s GitHub (https://github.com/stegiopast/ClearFinder). We provide instructions on how to set up the workflow using Conda environments. ClearFinder targets Linux systems and can access other operating systems using a virtual machine.

## Conclusions

ClearFinder offers a comprehensive and user-friendly solution tailored for scientists from various disciplines to process whole-brain light-sheet microscopy imaging data efficiently. Key features include:

- **Intuitive interface:** ClearFinder provides a guided platform for detecting and assigning cells throughout the entire mouse brain volume using the capabilities of the ClearMap and CellFinder tools
- **Robust installation process:** by maintaining independent virtual environments, ClearFinder ensures a robust and standardized installation process, simplifying the setup for users
- **Enhanced functionality:** ClearFinder extends basic processing capabilities by incorporating additional features such as result visualization through heatmap plotting and basic statistical analysis including PCA and box plots.

Moreover, using ClearFinder to cross-analyze a dataset, we demonstrated discrepancies in cell detection performance, raising the reproducibility issue within the field. Our tool allows the researchers to visually inspect the quality of cell detection, visualize the results and focus on consistent data from both analysis pipelines.

## Availability and requirements

**Project name:** ClearFinder

**Project home page:** https://github.com/stegiopast/ClearFinder

**Operating system(s):** Linux

**Programming language:** Python

**Other requirements:** Processors: 20 (Threats), RAM: 128GB

**License:** GNU General Public License v3.0

### List of abbreviations

DBE: Dibenzyl ether
DCM: Dichloromethane
DMSO: Dimethyl sulfoxide
GFP: Green fluorescent protein
GUI: Graphical user interface
i.p.: Intraperitoneal
IEG: Immediate-early genes
NDS: Normal donkey serum
PBS: Phosphate-buffered saline
PCA: Principal component analysis
PFA: Paraformaldehyde
PTx0.5: PBS with 0.5% TritonX-100
RI: Refractive index
RT: Room temperature

## Declarations

### Ethics approval and consent to participate

All experiments were performed according to the European Community’s Council Directive of 22 September 2010 (2010/63EU) and approved by the respective agency of the State Rhineland-Palatinate (Landesuntersuchungsamt, permit number G-17-1-021)

## Availability of data and materials

Data available upon request

## Authors’ contributions

Data analysis and code writing: SP with the help of SS, PH, NM. Data acquisition and preprocessing: MT, MC and IK. Writing – Original Draft Preparation, MT and IK. Writing – Review & Editing: MT, IK, BL, SG. Funding Acquisition: MT, BL, SG. Project Administration: MT. Supervision: MT and SG. All authors read and approved the final manuscript.

## Acknowledgements

We would like to thank Vladyslav Tsilytskyi for his help in proof-reading the code.

## Supplementary materials

1. Embedded ontology files for #1, #2, #3 analyzed with ClearMap and CellFinder
2. Parameters of cell detection used for analysis of data
3. Suppl. Fig. 1 Cell detection efficiency in cortical regions.
4. Suppl.Fig. 2 Cell detection efficiency in subcortical regions.

**Supplementary Figure 1.**
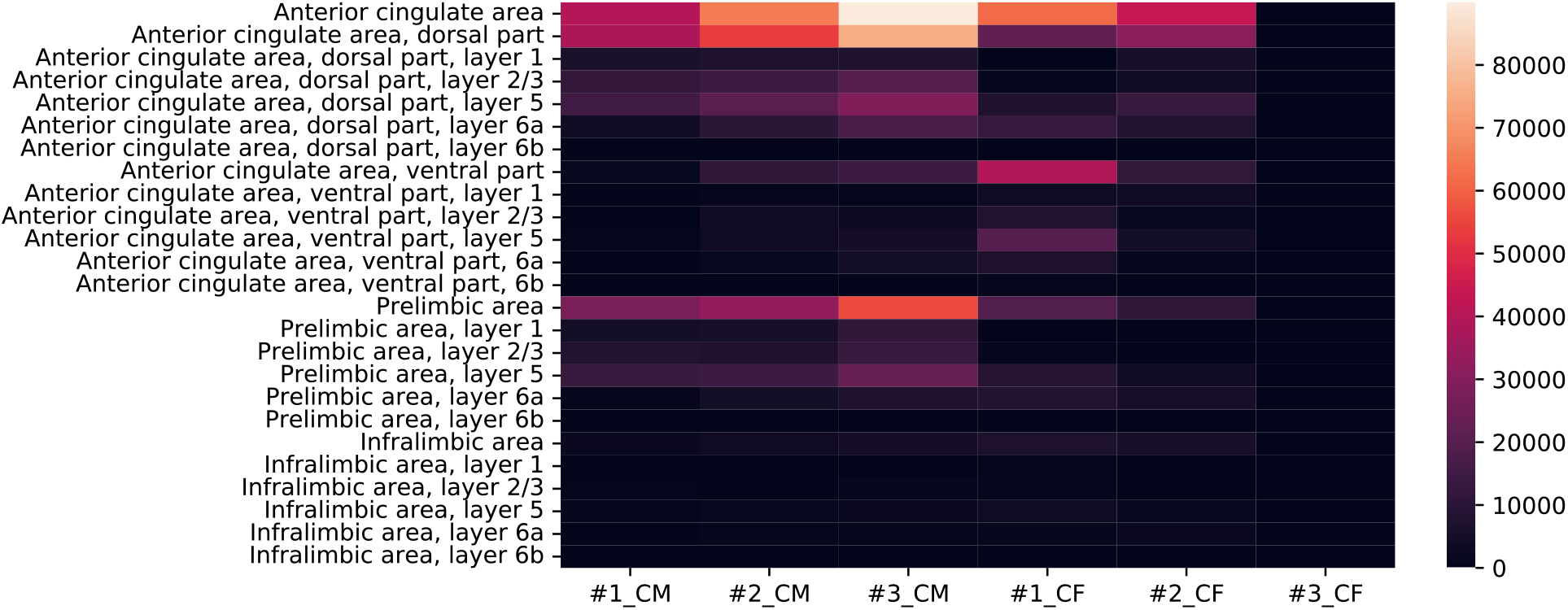
Cell detection efficiency in cortical regions. Heatmap indicating the number of cells detected in the subregions of the anterior cingulate, prelimbic, and infralimbic areas as detected in each sample by ClearMap (CM) and CellFinder (CF) using the same threshold. Color-bar corresponds to cell count. Regions with no cell counts were not accounted in the heatmap.

**Supplementary Figure 2.**
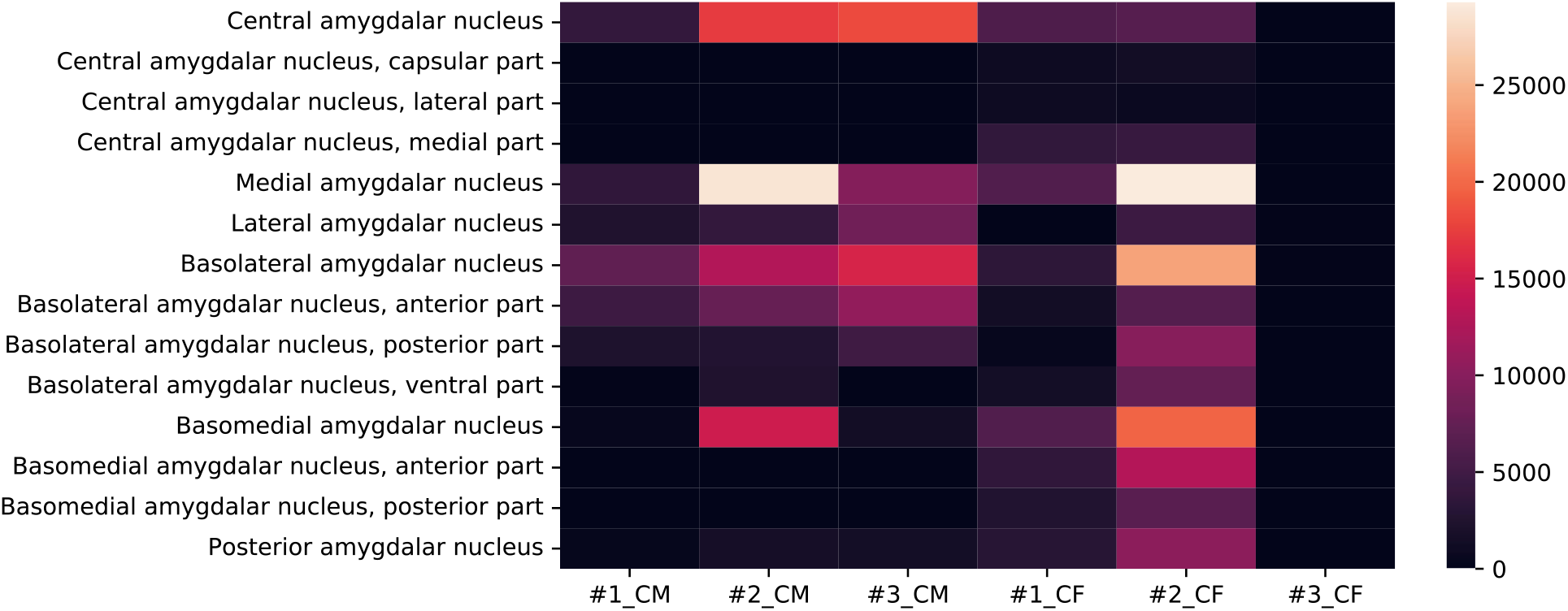
Cell detection efficiency in subcortical regions. Heatmap indicating the number of cells detected in the subregions of the central, medial, lateral, basolateral, basomedial, and posterior amygdalar nuclei as detected in each sample by ClearMap (CM) and CellFinder (CF) using the same threshold. Color-bar corresponds to cell count. Regions with no cell counts were not accounted in the heatmap.

